# Differences in dynamic motor selection in stuttering

**DOI:** 10.64898/2026.02.14.705922

**Authors:** Irene Echeverria-Altuna, Birtan Demirel, Sage E.P. Boettcher, Kate E. Watkins, Anna C. Nobre

## Abstract

Stuttering involves interruptions to the smooth flow of speech occurring mostly at syllable onset.^1^ Speech fluency is enhanced in people who stutter (PWS) by external timing cues.^2^ This has been taken to indicate that difficulties in the temporal organisation of action selection and initiation during speech contribute to stuttering.^3^ An important unanswered question is whether putative temporal coordination difficulties are specific to speech or generalise to other actions. Here, we examined the temporal organisation of hand action selection in PWS. Twenty PWS and twenty typically fluent speakers (TFS) underwent magnetoencephalography (MEG) recording while performing a visuomotor working-memory task that encouraged temporally specific selection, preparation, and shifts between hand actions. Lateralised sensorimotor mu/beta-frequency (8-30 Hz) activity modulation accompanying hand-action prioritisation was weaker in PWS than TFS. Strikingly, this effect was specific to a period of high uncertainty regarding which action to select and when. Despite these differences, lateralised mu/beta modulation was functionally related to reaction times in both groups and reaction times were well matched between PWS and TFS. The findings suggest a general disruption of temporal structuring of action selection and preparation in stuttering.

## Results

Stuttered speech is characterised by audible repetitions and prolongations of speech sounds, and inaudible blocks or pauses that interrupt the flow of speech. These disfluencies occur mostly at syllable onset.^1^ Speech fluency can be enhanced in people who stutter (PWS) by providing external rhythmic cues, such as a metronome, to time speech production.^2^ Thus, stuttering has been related to difficulties in timing and initiation of speech and the underlying neural circuits.^1,3,4^

Lateralised changes in sensorimotor mu/beta-frequency (8-30 Hz) activity are related to action selection and preparation.^5,6^ Previous magnetoencephalography (MEG) and electroencephalography (EEG) studies in stuttering have revealed differences in sensorimotor mu/beta-frequency (8-30 Hz) activity before and during speech production.^7–13^ Specifically, attenuation of beta-frequency (13-30 Hz) activity in preparation for speaking is enhanced in PWS compared with typically fluent speakers^8^ (TFS), although this is not always the case.^10^ Moreover, beta power increases before stuttered compared with fluent utterances have been found in PWS.^13^

Emerging evidence highlights broader difficulties in the temporal structuring of non-speech behaviours in stuttering. For example, behavioural performance and beta-frequency (13-30 Hz) EEG activity during finger-tapping tasks is affected in children who stutter^14^. Moreover, PWS show differences in beta-frequency EEG activity during auditory processing of speech and rhythmic sound sequences.^15–17^

Given the proposed difficulties with the temporal organisation of speech in stuttering, an important open question is whether there is a more general anomaly in temporally structuring action selection and preparation. Such a difficulty may be more evident in speech behaviour given the challenging temporal organisation required for selecting, sequencing, and coordinating movements across many articulators. However, methods with high temporal granularity may reveal similar difficulties when selecting and preparing other, non-speech actions.

Here, PWS (n = 20) and TFS (n = 20) performed a well-validated visuomotor working-memory task involving temporally organised hand action selection and preparation while MEG was recorded.^18^ The task required reporting the orientation of one of two memorised tilted bars. Each bar was uniquely associated with a response hand (left or right) based on its tilt (left or right). In most trials, two events predicted which bar would be probed (and the associated response hand): an informative retro-cue and the duration of the delay between the retro-cue and the probe. Together, the retro-cue and internal monitoring of delay duration encouraged self-guided and temporally structured prioritisation of different hand actions over time (**Figure 2a**).^18^ The PWS and TFS groups were well matched for handedness, gender, ethnicity, and education level (**Table S1**).

### PWS and TFS used retro-cues and delay duration to proactively prepare responses

Reaction time (RT) data were analysed using analysis of variance (ANOVA) testing the main effects of **cue** informativeness (informative vs. noninformative), **delay** duration (short vs. long), and **group** (PWS vs. TFS) and their interactions (**Figure 1b-c**). Responses were faster following informative vs. noninformative cues (F(*1,38*) = 139.39, ***p < .001, **η**^2^ = .34), and for long vs. short delays (F(*1,38*) = 149.41, ***p < .001, **η**^2^ = .2). The benefit of informative cues was greatest at long delays (interaction between cue and delay: F(*1,38*) = 13.43, ***p < .001, **η**^2^ = .02; post-hoc t-tests: short, t(*77.6*) = −2.18, *p = .03, d = .49; long, t(*74.2*) = −7.32, ***p < .001, d = 1.64). Response speed of PWS and TFS did not differ and there were no other significant interactions (**Table S2**). The benefit of informative cues on RT across both delays showed that participants used retro-cues and monitored delay duration to prepare their responses proactively.

**Figure 1.**
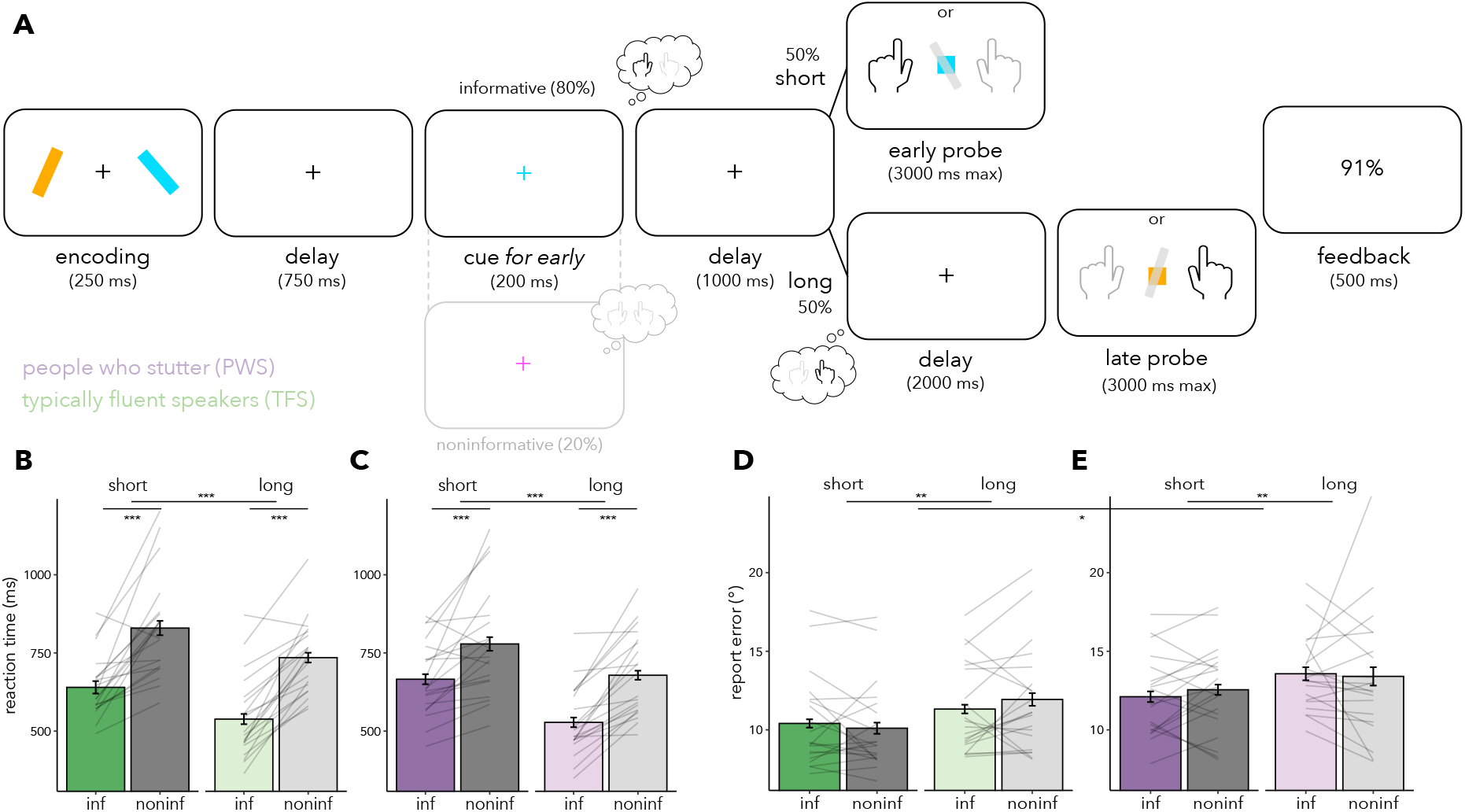
Task design and behavioural results in typically fluent speakers (TFS) and people who stutter (PWS). (a) Trial schematic. Two coloured, tilted bars (on the left and right) were displayed (one tilted to the left and the other to the right). Participants were probed to report the orientation of one of the encoded bars at the end of each trial. In 80% of trials (informative), a retro-cue matching the colour of one of the two bars was displayed; in noninformative trials (20%), a cue with a different colour appeared. In informative trials, participants reported the cued item if probed after a short delay (1s) or reported the other item if probed after a long delay (3s). In noninformative trials, the item to report was unpredictable if probed after a short or long delay. Participants received trial-wise feedback about their response accuracy as a percentage. (b-c) Mean reaction times (RT) in informative (coloured) and noninformative (grey) correct trials at short (dark) and long (light) delays for TFS (b; green) and PWS (c; purple). (d-e) Average absolute report error (°) in informative (coloured) and noninformative (grey) correct trials at short (dark) and long (light) delays for TFS (d; green) and PWS (e; purple). Thin grey lines represent individual participants, and error bars depict the standard error of the mean (SEM). Asterisks represent statistical significance (*p < .05; **p < .01; ***p <.001). See also Tables S1 and S2.

Both groups performed the task highly accurately, with average report errors below 15°. An ANOVA on report error revealed that TFS were, on average, more accurate than PWS in their orientation reports (F(*1,38*) = 6.52, *p = .02, **η**^2^ = .08). Both groups were more accurate after a short delay than a long delay (F(*1,38*) = 7.48, **p = .01, **η**^2^ = .03; **Figure 1d-e**). No other significant effects or interactions were observed (**Table S2**). Thus, task performance, as measured with report error, across all conditions was slightly less accurate in PWS compared to TFS.

### Time-specific differences in lateralised sensorimotor mu/beta activity modulation

The use of temporal predictions to prioritise and shift between left- and right-hand actions is mirrored by dynamic changes in lateralised sensorimotor mu/beta (8-30 Hz) activity.^18^ Lateralised mu/beta modulation in motor areas contralateral vs. ipsilateral to hand actions provides a functional index of the relative selection of one action amongst two competing hand actions.^5,19^ We built on this to investigate putative differences in the time course of selective hand action prioritisation between PWS and TFS.

First, we contrasted lateralised time-frequency activity in M1 parcels (see **STAR Methods** for details) locked to retro-cue onset based on the left vs. right hand action linked to the cued item. In TFS, cluster-based permutation testing of lateralised time-frequency activity revealed a pronounced modulation of mu/beta activity that followed hand action selection and preparation. Mu/beta power was initially relatively diminished contralaterally (vs. ipsilaterally) to the cued action followed by a shift toward diminished activity contralaterally to the other action after the first delay (**Figure 2a**; from left to right – first cluster: ***p < .001; time range: .59-1.63 s, frequency range: 2-36 Hz; second cluster: ***p < .001; time range: 1.8-3.2 s, frequency range: 7-37 Hz). In contrast, PWS had no detectable modulation of time-frequency activity in M1 contralateral-vs-ipsilateral to the cued action (no clusters found), followed by a pronounced mu/beta modulation contra-vs-ipsi to the action associated with the opposite (uncued) bar (**Figure 2b**; from left to right – first cluster:*p = .04; time range: 1.79-2.2 s, frequency range: 18-30 Hz; second cluster: **p =.005; time range: 2.35-2.81 s, frequency range: 2-36 Hz; third cluster: **p = .009; time range: 2.87-3.2 s, frequency range: 11-36 Hz).

**Figure 2.**
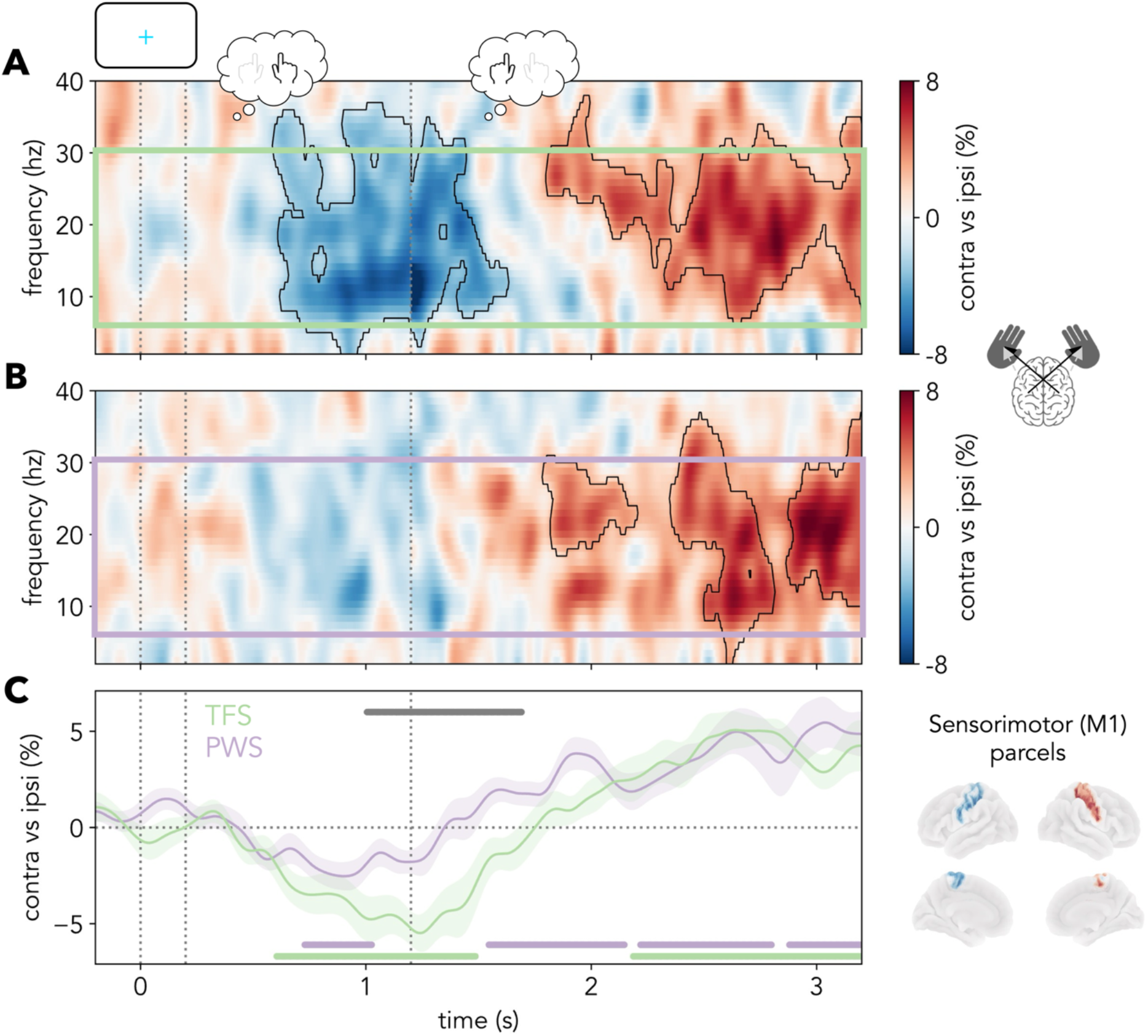
Lateralised time-frequency activity in sensorimotor (M1) parcels contralateral vs. ipsilateral to the cued action locked to cue onset. (a-b) Contrast between M1-localised MEG time-frequency-activity contralateral vs. ipsilateral to the cued action, divided by summed contralateral and ipsilateral activity (expressed as a percentage) in informative, correct trials in TFS (a; top) and PWS (b; middle). (c) Cross-participant average mu/beta (8-30 Hz) activity difference between M1 parcels contralateral and ipsilateral to the cued action in informative correct trials. The black outlines in the time-frequency spectra indicate statistically significant clusters. Shaded areas in c represent the SEM. Cluster-permutation corrected significant timepoints are indicated with horizontal lines in c. The first part (−0.2-1.2 s) of the time-frequency spectra in a and b and of the time course in c corresponds to the average of short and long delay trials, and the second part (1.2-3.2 s) corresponds to long delay trials only. The vertical dotted lines indicate (from left to right) the onset (0 s) and offset (0.2 s) of the retro-cue and the time of probe appearance in short delay trials (1.2 s). PWS: people who stutter; TFS: typically fluent speakers. See also Figures S1, S2 and S3 and Table S3.

Next, we contrasted the average mu/beta-band (8-30 Hz) modulation time courses between PWS and TFS (**Figure 2c**). When restricting the analyses to the mu/beta (8-30 Hz) band, both groups showed significant modulation of lateralised activity related to first- and second-action prioritisation (TFS: first cluster: **p = .002; time range: 0.6-1.49 s; second cluster: ***p < .001; time range: 2.2-3.2 s; PWS: first cluster: *p = .02; time range: 0.73-1.03 s; second cluster: **p = .008; time range: 1.5-2.14 s; third cluster: **p = .008; time range: 2.21-2.8 s; fourth cluster: *p = .03; time range: 2.88-3.2s). Strikingly, lateralised sensorimotor mu/beta modulation was significantly weaker in PWS than TFS when the predicted hand action shifted (cluster: **p = .008; time range: 1-1.69 s).

In contrast, no group differences were observed in the time course of lateralised modulation of occipital alpha (8-12 Hz) activity related to the sensory selection of item locations in working memory (**Figure S1**).^18^

To investigate the temporal dynamics of mu/beta activity underlying the observed differences, we explored the time course of mu/beta (8-30 Hz) power in sensorimotor areas contralateral and ipsilateral to the cued action (**Figure S2**). Interestingly, this seemed to point to a generally weaker modulation of mu/beta frequency activity across both hemispheres in PWS vs. TFS (**Figures S2a** and **S2b**). Nevertheless, this difference was not statistically significant. To separate the contributions of contralateral and ipsilateral areas to the observed group differences in lateralised mu/beta modulation specifically, we compared mu/beta power during the time window showing significant group differences in lateralised mu/beta modulation (**Figure 2c**; 1-1.7 s) in contralateral and ipsilateral sensorimotor parcels. This comparison revealed that mu/beta power attenuation in the hemisphere contralateral to the cued action was stronger in TFS than PWS (t(37.94) = 2.1, p = .04, d = .66; **Figure S2c**). No differences in mu/beta power were observed between groups in the hemisphere ipsilateral to the cued action during the same time window (t(37.96) = −.11, p = .91, d = .03; **Figure S2d**). Together, this suggests that the observed group differences in lateralised mu/beta modulation around shift time were driven by changes in sensorimotor cortices contralateral to the cued action.

### Differences in lateralised mu/beta modulation amplitude, not timing, in PWS

Visual inspection of the lateralised mu/beta modulation time courses (**Figure 2c**) revealed two main differences between PWS and TFS. First, the amplitude of lateralised mu/beta modulation was weaker in PWS around 1-1.7 s after informative cue onset. Secondly, the sign-reversal (zero-crossing) of the lateralised mu/beta modulation time course seemed to happen earlier in PWS (average time at zero-crossing: 1.35 s) than in TFS (average time at zero-crossing: 1.75 s). This may reflect true group differences in hand-action shift time (i.e., earlier shifts or higher trial-wise variability in shift time in PWS). Alternatively, it may result from differences in the amplitude of lateralised mu/beta modulation.

To disambiguate these possibilities (i.e., amplitude- vs. timing-related differences in lateralised mu/beta modulation vs. both), we used a leave-one-subject-out lagged template matching approach to independently estimate single-trial values of sensorimotor mu/beta shift time and amplitude (see **STAR Methods**). Linear modelling of trial-wise shift time while controlling for amplitude revealed that PWS and TFS did not differ in the sign-reversal time (shift time) of the lateralised mu/beta activity modulation time course (**Table S3**). Other supplementary analyses revealed no group differences in MEG signal-to-noise ratio, including parcel-wise power spectra (**Figure S3a-c**), trial-wise standard deviation of lateralised mu/beta modulation time series and inter-trial phase coherence of M1 activity (see **STAR Methods**).

### Lateralised sensorimotor mu/beta modulation is linked to task performance

Finally, we tested whether lateralised mu/beta activity modulation was linked to task performance. Trials for individual participants were divided into *fast* or *slow* based on the participant-wise median RT. Fast trials were associated with stronger lateralised sensorimotor mu/beta modulation compared with slow trials in TFS (**Figure 3a,c**; cluster: **p = .003; time range: 2.2-3.2 s; short delay: t(*35.62*) = −2.18, p = .09, d = .55; long delay: t(*36.68*) = 3.51, **p = .001, d = 1.11).^18^ The same pattern was observed in PWS (**Figure 3b,d**; cluster: *p = .04; time range: 2.09-2.54 s; short delay: t(*37.92*) = −2.85, **p = .007, d = .9; long delay: t(*25.96*) = −2.09, *p = .04 d =.66). Interestingly, the association between faster responses and stronger lateralised mu/beta modulation in PWS held even in the short delay, despite overall weaker modulation of mu/beta activity (**Figure 2c**). Thus, lateralised sensorimotor mu/beta modulation seemed to be functionally related to hand action prioritisation in both groups.

**Figure 3.**
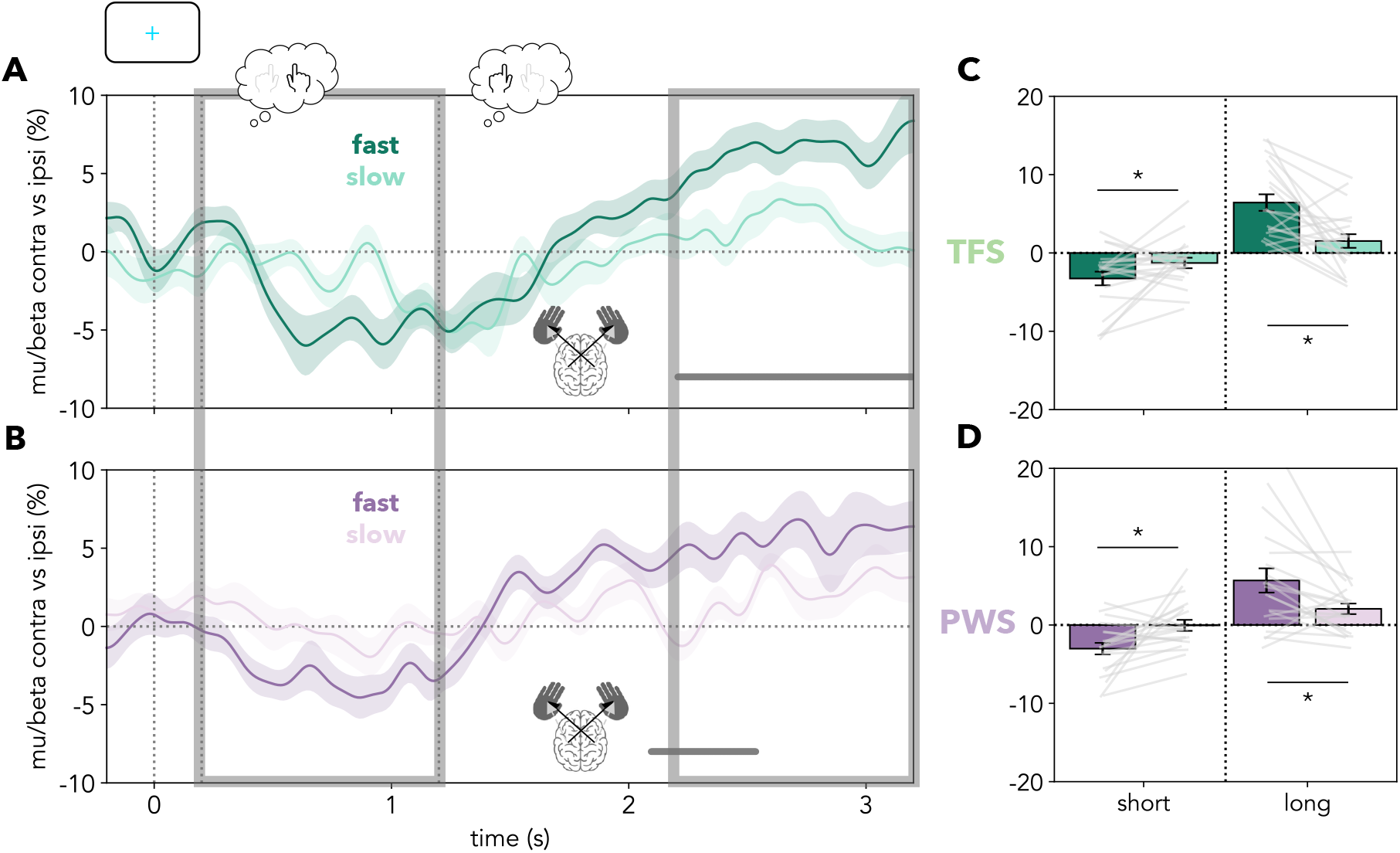
Relation between lateralised sensorimotor mu/beta activity modulation and RT in typically fluent speakers (TFS) and people who stutter (PWS). (a-b) Sensorimotor mu/beta (8-30 Hz) activity contralateral vs. ipsilateral (expressed as a percentage) to the cued hand-action in trials categorised as fast (dark) vs. slow (light) based on participant-wise median splits of RT in TFS (a; top; green) and PWS (b; bottom; purple). (c-d) Bar plot of average lateralised mu/beta modulation in the early (0.2-1.2 s) and late (2.2-3.2 s) time windows, depicted by the shaded grey rectangles in a and b. Coloured shaded areas represent the SEM, and cluster-permutation corrected significant timepoints are indicated with horizontal lines in a and b. The first part (−0.2-1.2 s) of the time courses in a and b corresponds to the average of short and long delay trials and the second part (1.2-3.2 s) corresponds to long delay trials only. The vertical dotted lines represent (from left to right) the onset (0 s) and offset (0.2 s) of the retro-cue and the time of probe appearance in short delay trials (1.2 s). Thin grey lines represent individual participants, and error bars depict the SEM. Asterisks represent statistical significance (*p < .05; **p < .01; ***p < .001). See also Figure S3 and Table S3.

Finally, despite being generally weaker, the magnitude of lateralised sensorimotor mu/beta activity modulation around between-action shift time (1-1.7 s after the cue) in PWS did not correlate with stuttering severity (as measured with the SSI-4^20^), stuttering experience (as indexed by OASES^21^ scores), or anticipatory awareness of stuttering (as indicated by PAiS^22^ results; **Figure S3d-f**).

## Discussion

Using a visuomotor working-memory task that encourages using the passage of time to shift dynamically between actions, we discovered that stuttering is associated with differences in how temporal structures guide the selection and preparation of hand actions. Specifically, lateralised sensorimotor mu/beta-frequency (8-30 Hz) activity modulation related to selective hand action prioritisation was weaker in PWS than TFS.

This difference in lateralised mu/beta modulation was temporally specific. The period of maximal difference between PWS and TFS coincided with the expected time of shift between hand actions. This period is characterised by ambiguity regarding which response (left- or right-hand action) will be required and when that response will need to be initiated. In contrast, no group differences in lateralised sensorimotor mu/beta modulation were observed later in the delay period, when the to-be-probed hand action and its timing were predictable. It is also worth noting that mu/beta activity modulation in motor areas contralateral and ipsilateral to the cued action seemed to be generally weaker in PWS than in TFS, though this difference was not statistically significant. Together, these findings indicate that neural activity differences related to stuttering may emerge during temporal structuring of the selection and preparation of multiple, potentially competing, action plans. More broadly, the present study underscores the importance of using tasks with temporal structuring demands and methods with high temporal resolution for elucidating the cognitive and neural processes underlying stuttering.

Together, these results suggest that stuttering involves differences in the temporal structuring of action selection and preparation in a way that is not specific to speech.^3,23^ Speculatively, the primary manifestation of stuttering symptoms in speech may arise from the greater demands for fine-grained temporal structuring required for phoneme selection, sequencing and articulation.

The present study extends existing literature reporting differences in speech-related mu/beta activity in stuttering^7–10,13,16,17^ by additionally highlighting differences in mu/beta activity related to non-speech action selection and preparation in adults who stutter. The functional significance of lateralised sensorimotor mu/beta-frequency activity modulations remains debated.^24^ Here, we interpret modulations in mu/beta-frequency activity as signalling changes in neural excitability in the sensorimotor system, relating to action selection, prioritisation, preparation, and initiation.^6^ Importantly, relative modulations of mu/beta power in sensorimotor areas contralateral vs. ipsilateral to hand actions reflect the selective prioritisation of one action relative to other, competing actions.^5,19^

Interestingly, several studies have linked changes in mu/beta activity to the engagement of cortico–basal ganglia–thalamo–cortical loops,^24^ in which abnormal function has also been implicated in stuttering.^3,23^ Specifically, neuroimaging studies have revealed structural and functional differences between PWS and TFS in neural systems responsible for selecting, initiating, and executing action sequences (i.e., speech), including basal ganglia, cerebellum, supplementary motor area (SMA), and inferior frontal gyrus (IFG).^25,26^ The magnitude of the contingent negative variation (CNV) potential, related to the engagement of cortico-basal ganglia-thalamo-cortical loops, also seems to differ between PWS and TFS prior to speech production.^27^ Interestingly, transcranial magnetic stimulation (TMS) of the SMA in PWS and TFS revealed changes in the magnitude and timing of TMS-evoked EEG activity in the SMA and left IFG, further pointing to differences in the same cortico-subcortical network.^28^ Notably, areas within these networks were differentially engaged in PWS compared with TFS during the preparatory stage of a go/no-go task.^29^ Although the contribution of cortico–basal ganglia–thalamo– cortical networks to the differences in lateralised mu/beta activity observed in this task is difficult to establish with MEG, the present findings are broadly consistent with studies reporting differences in neural systems related to the temporal structuring of actions in PWS and to existing computational models of stuttering.^4^

The group differences in lateralised mu/beta modulation appeared to result from time-specific changes in the amplitude of such modulation, especially in contralateral sensorimotor cortices, rather than from differences in between-action shift time or shift-time variability. Thus, the observed effects are unlikely to reflect differences in neural signals specifying when to shift between actions and might instead reflect anomalies in other dimensions of motor competition, selection, or preparation. These may include unstable forward modelling,^10^ inefficient action selection and preparation,^7,30^ altered inhibitory motor control,^13^ and/or differences in action timing or temporal processing.^3,15,23^ Further research will be instrumental to delineate the specific cognitive and neural mechanisms underlying the observed differences in sensorimotor mu/beta activity in stuttering.

Strikingly, lateralised mu/beta modulation remained functionally related to behaviour despite being weakened in PWS (**Figure 3**). Moreover, response times were comparable between PWS and TFS at both the short and long delays, despite marked group differences in lateralised sensorimotor mu/beta modulation between delays (**Figure 2c**). The observation that neural activity related to action selection differed despite intact behavioural performance may point to the engagement of alternative, potentially compensatory, neural processes in PWS that preserve the speed of manual responses in this task.

The observed differences between PWS and TFS were specific to lateralised sensorimotor mu/beta modulation related to dynamic hand action selection. In contrast, PWS did not show differences in the time course of lateralised occipital alpha-frequency (8–12 Hz) activity associated with the dynamic prioritisation of visuospatial working-memory contents.^18,19^ This dissociation suggests that stuttering may be associated with differences in the temporal selection and preparation of motor, but not visuospatial, contents. Whether the temporally structured selection of auditory or verbal working-memory contents is affected is an interesting avenue for future research.

In conclusion, the present study found differences in lateralised sensorimotor mu/beta activity during temporally structured shifts between hand actions in PWS. Thus, stuttering may involve alterations in the temporal structuring of action selection and preparation beyond speech. The observed differences were specific to moments that required structuring several possible actions and action initiation times, a feature of this task which is shared with speech. More broadly, the present results highlight the importance of experimental designs and analytical frameworks with high temporal and functional granularity for isolating the cognitive and neural mechanisms that contribute to stuttering.

## Acknowledgements

The authors would like to thank Amelia Rock, Anne Schroder, and the teams responsible for the Oxford Centre for Human Brain Activity (OHBA) MRI and MEG facilities for their invaluable assistance with data collection. The authors also thank the members of the Brain and Cognition Lab for helpful discussions, and all participants who generously contributed their time and effort to this study.

This research was funded by a Wellcome Trust PhD Studentship (102170/Z/13/Z), a Rafael del Pino Excellence Studentship and a Programa Posdoctoral de Perfeccionamiento de Personal Investigador Doctor del Gobierno Vasco fellowship to I.E.A.; and a James S. McDonnell Foundation Understanding Human Cognition Collaborative Award (220020448) and a Wellcome Trust Senior Investigator Award (104571/Z/14/Z) to A.C.N. This research was also supported by the Dominic Barker Trust (Registered Charity no. 1063491). The Wellcome Centre for Integrative Neuroimaging is supported by core funding from the Wellcome Trust (203139/Z/16/Z and 203139/A/16/Z). This research/study/project is funded/supported by the NIHR Oxford Health Biomedical Research Centre (NIHR203316). This research is supported by the Basque Government through the BERC 2022-2025 program and Funded by the Spanish State Research Agency through BCBL Severo Ochoa excellence accreditation CEX2020-001010/AEI/10.13039/501100011033.

For the purpose of open access, the author has applied a CCBY public copyright license to any Autor Accepted Manuscript version arising from this submission.

## Author contributions

Conceptualization: I.E.A., S.E.P.B., K.E.W., and A.C.N. Methodology: I.E.A., S.E.P.B., and A.C.N. Software and Formal Analysis: I.E.A. Investigation: I.E.A. and B.D. Resources: K.E.W., and A.C.N. Data Curation: I.E.A., B.D., and K.E.W. Writing – original draft: I.E.A., K.E.W., and A.C.N. Writing – review and editing: all authors. Visualization: I.E.A., S.E.P.B., K.E.W., and A.C.N. Supervision: K.E.W. and A.C.N. Project administration: I.E.A., K.E.W., and A.C.N. Funding Acquisition: I.E.A., K.E.W., and A.C.N.

