## Supplementary Materials for "Differences in dynamic motor selection in stuttering"

### STAR Methods

#### Experimental model and study participant details

The study protocol was approved by the Central University Research Ethics Committee of the University of Oxford (R51267/RE004). Twenty people who stutter (PWS) and twenty age-, gender-, ethnicity-, and education-matched typically fluent speakers (TFS) participated in this study (**Table S1**). Informed consent was obtained before the study according to the Declaration of Helsinki, and participants were reimbursed for participation (£20/h). Participants were screened to ensure suitability for MEG and magnetic resonance imaging (MRI). All participants were native English speakers, had no history of neurological or psychiatric disorders, and self-reported to have normal or corrected-to-normal vision and hearing.

Speaking and reading samples were recorded using video and audio in PWS for off-line assessment using the Stuttering Severity Index-4 (SSI-4).[20] PWS also completed the Overall Assessment of the Speaker's Experience of Stuttering questionnaire (OASES)[21] and the Premonitory Awareness in Stuttering Scale (PAiS).[22]

#### Method details

##### Task and procedure

Participants performed a visuomotor working-memory task designed to study the selective and dynamic prioritisation of working-memory items and hand actions (for detailed task description see [18]; **Figure 1a**). Stimuli were presented using the Psychophysics Toolbox version 3.0.18[31] using Matlab 2022a[32] on a translucent whiteboard with an LED projector (ProPixx, VPixx Technologies Inc., Saint-Bruno, Quebec, Canada) placed 120 cm away from the participant (refresh rate: 120 Hz; projection area: 55 x 51 cm; resolution: 1920 x 1080 pixel). Manual responses were recorded using two bimanual fibre-optic response devices, one for each hand.

The task procedure mostly followed a previous study.[18] In each trial, two coloured, tilted bars were displayed, one on the left and one on the right side of the screen (bar width: 38 pixels; height: 192 pixels; distance from screen centre: 5.2 DVA). One bar was tilted leftward and the other rightward with respect to the vertical axis. Across trials, the location and tilt of the bars were orthogonal.[19] At the end of each trial, a coloured square appeared (response probe; width: 38 pixels), prompting participants to report the tilt of the bar with the matching colour from memory. Importantly, the tilt of each bar (left vs. right) was directly linked to the hand participants should use to respond (left vs. right). Thus, each working-memory item was uniquely linked to a manual action.[19]

In most trials (80%; informative), the colour of the central fixation cross changed (retro-cue)[33] to match the colour of one of the memorised bars, thus prompting participants to select one working-memory item, and the associated response hand, in anticipation of the response probe. In half of the informative trials, the response probe appeared 1 s after retro-cue offset (short delay trials). In the other half, the response probe appeared 3 s after cue offset (long delay trials; **Figure 1a**).

Crucially, in informative trials, the two pieces of information predicted the item to be probed with 100% validity: 1) the colour of the retro-cue; 2) the duration of the delay between the retro-cue and the response probe. When the delay was short, participants were always probed about the bar with the colour matching the retro-cue. Alternatively, in long delay trials, participants were always probed about the other (uncued) bar.

In summary, based on the colour of the retro-cue, participants could anticipate the item to be probed and the associated hand action. They should initially focus internal attention on the cued item and prepare the associated response hand. In long trials, once the short interval lapsed, participants should shift their focus to the other item and action. Importantly, the shift from prioritising one item and action to the other was self-guided and depended on the internal monitoring of delay duration.[18]

In a small proportion of trials (20%, noninformative), the colour of the retro-cue did not match the colour of either bar. In these trials, participants were unable to anticipate the to-be-probed item and associated response at either delay duration.

After the probe appeared, participants reported the orientation of the encoded bar with the matching colour by pressing and holding one of two keys with the left or right hand, which caused a central grey bar to rotate leftwards or rightwards, respectively.[18] When the grey bar reached the desired orientation, participants released the key thus confirming their response.

After each trial, participants received feedback about the deviation of their report from the probed bar's orientation. Participants performed four task blocks with 32 trials each (~6 mins/block). In each block, trials with the various experimental factors were randomly interspersed: bar location, bar tilt (related to response hand), retro-cue colour, retro-cue informativeness (informative vs. noninformative), and interval duration (short vs. long).[18]

#### MEG: acquisition and preprocessing

Whole-head MEG recordings were acquired using a 306-channel system (204 first-order planar gradiometers, 102 magnetometers; TRIUX neo, MEGIN OY, Espoo, Finland). Data were recorded with a sampling rate of 1000 Hz and an online filter between 0.03 and 300 Hz. Participants were seated during the task and head position in the scanner was tracked (see [34] for details). Electrocardiography (ECG) and vertical and horizontal electrooculography (EOG) were measured. Participants were instructed to avoid moving and to keep their gaze fixed on the central cross during the task. Structural MRI T1-weighted scans were acquired at the Oxford Centre for Human Brain Activity using a Siemens Prisma 3T scanner.

Spatiotemporal signal space separation and movement compensation of the raw MEG data were performed using Elekta MaxFilter version 2.2 (Neuromag). MEG recordings (two per participant) were aligned to the head position of the first recording. All resulting MEG data were preprocessed and analysed in Python 3.11.9[35] using MNE-Python[36] (version 1.5.1), osl-ephys[37] and custom-made scripts. First, data were high-pass filtered at 0.05 Hz, low-pass filtered at 125 Hz, and notch-filtered (50 Hz). Noisy channels were identified using a generalised extreme studentised deviate (ESD) test[38] at a 0.05 significance threshold and interpolated using spline interpolation. Independent component analysis (ICA) identified "artifactual" eye- and heart-related activity based on correlations between individual ICs and EOG and ECG recordings, as well as visual inspection of IC time courses and topographies. An average of 3.19 ICs (SD: 0.82; range: 2-5) were subtracted. There were no differences in the number of "artifactual" ICs between the groups.

#### MEG: source localisation

Sensor-level MEG data were source-reconstructed using osl-ephys.[37] First, MRI and MEG coordinate systems were co-registered by matching the digitised anatomical landmarks and scalp shape points onto participants' T1 scans. A single-layer (inner skull) forward model was computed, and source activity was estimated on an 8 × 8 × 8 mm volumetric grid using a unit-noise-gain invariant Linearly Constrained Minimum Variance (LCMV) beamformer.[39] The beamformer weights were computed from the leadfields of the forward model, a sensor noise covariance matrix, and a data covariance matrix. The noise covariance was modelled as diagonal, with entries given by the mean temporal variance for each sensor type (magnetometers and gradiometers). The data covariance matrix was estimated from the continuous preprocessed data, and regularisation was performed using Principal Component Analysis (PCA) to a rank of 60. This regularisation improves the robustness of the beamformer and is consistent with the default osl-ephys pipeline.[37]

Source activity was estimated at each grid point, and grid-wise time series were then parcellated into 52 anatomically defined cortical regions using an approximately symmetric atlas. This parcellation provides a standard dimensionality reduction for whole-brain analyses and ensures that the number of parcels does not exceed the effective data rank, thereby avoiding rank deficiency in subsequent analyses.[40] Parcel time courses were computed by applying PCA to the demeaned grid time series within each region and retaining the first principal component to provide a sign-consistent representative signal.

### Quantification and statistical analysis

#### Behavioural data analysis

Behavioural data were analysed using R[41] (version 4.2.1) and Rstudio[42] (version 2022.07.1). Reaction time (RT) was calculated as the time from probe onset until response initiation in correct trials. Report error was defined as the absolute difference in degrees (°) between the reported orientation and the orientation of the probed bar in trials where the correct hand was used to respond. Trials lacking a response, trials with RTs faster than 100 ms, trials with RTs slower than three times the SD of the average RT per participant, and trials with report errors higher than three times the SD of the participant-averaged report error were excluded from further analyses. A total of 6.35% (SD: 5.12%) of trials were rejected across participants.

The statistical significance of RT and report error across conditions and groups was tested using mixed-design analysis of variance (ANOVA) with cue informativeness (informative vs. noninformative) and delay duration (short vs. long) as within-subject factors, and group (PWS vs. TFS) as a between-subject factor. Given their skewed distributions, RT and report error were log-transformed before statistical testing. Post-hoc t-tests were used to test the direction of interaction effects.

#### MEG: epoching and time-frequency analyses

Parcel-wise time courses were epoched based on retro-cue onset (-0.25 s to 1.25 s in short trials and -0.25 s to 3.25 s in long trials). Activity during the baseline period

(-0.25 to 0 s) was subtracted from each epoch. Only informative "correct" trials (see above) were included in the MEG analyses.

The epoched parcel time courses were convolved into their time-frequency decompositions with Morlet wavelets with frequency-dependent widths from 2 to 40 Hz in steps of 1 Hz. The 50 ms around the edges of each epoch were cropped to remove edge artifacts related to the time–frequency decomposition. Lateralised mu/beta activity (8-30 Hz) modulation source localised to primary motor cortex (M1)[6,19,43] is a proxy of hand-action prioritization. Therefore, we estimated time-frequency activity in the parcels encompassing the left and right superior and inferior sensorimotor cortices[40] (hereon M1; **Figure 2c**). Time-frequency activity was calculated for left and right M1 separately when the corresponding right- and left-hand action was cued. The normalised difference in contralateral vs. ipsilateral activity was derived for each M1 parcel and then combined across hemispheres to derive an overall lateralisation index: [((contra – ipsi) / (contra + ipsi)) * 100]. Next, the contrasts across both sides were collapsed and averaged across participants. The average contra-vs-ipsi mu/beta activity modulation contrast was calculated per participant by averaging across the 8-30 Hz frequency band and smoothing the time courses using a Gaussian kernel with a standard deviation (SD) of 40 ms.

Lateralised alpha (8-12 Hz) activity modulation related to visuomotor working-memory content prioritisation was estimated in parcels encompassing primary and secondary visual cortices[19] using the procedure detailed above (**Figure S1**).[40]

The separate contributions of contralateral and ipsilateral sensorimotor cortices to the observed contra-vs-ipsi mu/beta modulations were assessed as follows. For each participant, baseline mu/beta (8-30 Hz) power (-0.2-0 s) was subtracted from the cue-locked mu/beta power time course and averaged over the time window showing a contra-vs-ipsi mu/beta modulation difference between groups (1-1.7 s; **Figure 2c**). The separate contralateral and ipsilateral measures were compared between groups with cluster-based premutation testing (for the time courses; see below) and independent-samples t-tests (for the participant averages during the time window of interest; **Figure S2**).

Cluster-based non-parametric permutation was used to evaluate MEG differences between conditions and groups. The contra-vs-ipsi time-frequency spectra were tested against a null hypothesis of 0 using one-sample cluster-based permutation testing.[44] The average time courses of lateralised alpha and mu/beta modulation between groups were tested using independent-sample cluster-based permutation testing.

#### MEG: relation with behaviour

For each participant, we identified trials with RTs faster or slower than their median RT, and action-related contra-vs-ipsi time-frequency spectra were calculated separately for faster-than-median and slower-than-median trials. Lateralised mu/beta activity modulation time-courses were contrasted between fast and slow trials using one-sample cluster-based permutation testing.

#### MEG: trial-by-trial lateralised mu/beta activity modulation analyses

Single-trial contra-vs-ipsi time-frequency spectra were estimated using Morlet wavelets for left- and right-action cue trials separately as detailed above. Then, time-frequency spectra from left- and right-action cue trials were concatenated, and the number of trials was equalised between participants through random downsampling.

A leave-one-subject-out (LOSO) lagged template matching approach was used to quantify potential differences between PWS and TFS in the amplitude and temporal alignment of single-trial lateralised mu/beta modulation time courses. Trial-wise mu/beta activity modulation was estimated by averaging the time-frequency spectra between 8-30 Hz and baseline (-0.2-0 s) z-scoring power from each trial. For each participant, a LOSO lateralised mu/beta modulation template was constructed by averaging the smoothed (40-ms Gaussian kernel) mu/beta modulation time courses of all participants except themselves. Each participant's smoothed single-trial modulation was then fitted to the corresponding LOSO template by minimizing the root-mean-square (RMS) error using the L-BFGS-B algorithm implemented in SciPy minimise:[45]

$$\text{trial}(t) \approx a * \text{template}(t - \tau) + b$$

where $\tau$ is the temporal shift (s), a is the amplitude scaling factor, and b is a baseline offset. Fits were initialised at $\tau = 0$ s, with a maximum allowable shift of ±2 s. Different values of maximum shift (0.5-2.5 s) confirmed the observed pattern of results.

This yielded trial-wise estimates of temporal shift ($\tau$) and amplitude (a). A linear mixed-effects model (LMM) as implemented in R with lmer[46] was then used to test for group (PWS vs. TFS) differences in $\tau$, while controlling for within- and between-participant changes in amplitude (a) and fit quality (RMS). The model was estimated using a maximum likelihood criterion, and the outputs were reported as unstandardised regression coefficients with t-statistics. Two-tailed tests and a 0.05 criterion for significance were used.

#### MEG: control analyses

We ensured that PWS and TFS were comparable in terms of data quality by estimating power spectra for each participant and parcel and comparing power spectra between groups using a general linear model (GLMspectra).[47] The variance (SD) of cue-locked lateralised mu/beta activity modulation over time was also calculated across trials and compared between PWS and TFS using cluster-based permutation testing. Finally, inter-trial phase coherence (ITPC) on cue-locked epoch-wise time-frequency activity was calculated using MNE-Python and averaged across participants from each group. Independent-samples cluster-based permutation testing compared ITPC between groups.

### Supplementary Figures and Tables

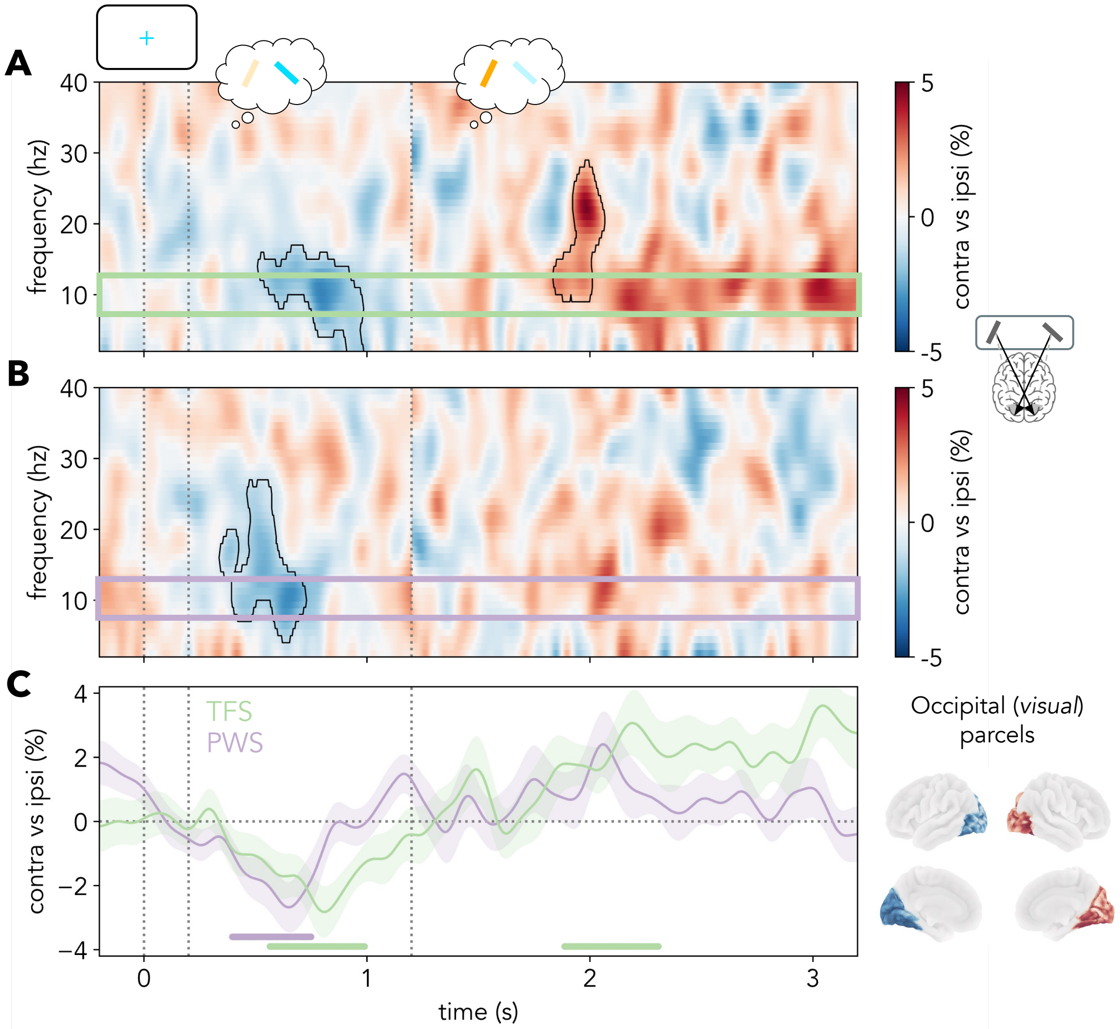


**Supplementary Figure 1. Lateralised time-frequency activity in visual parcels contralateral vs. ipsilateral to cued location, locked to cue onset. Related to Figure 2.** (a-b) Contrast between source-localised MEG time-frequency activity contralateral vs. ipsilateral to the cued location, divided by summed contralateral and ipsilateral activity (expressed as a percentage) in informative, correct trials in TFS (a; top) and PWS (b; middle). (c) Cross-participant average alpha (8-12 Hz) activity difference between visual parcels contralateral and ipsilateral to the cued location in informative correct trials. The black outlines in the time-frequency spectra indicate statistically significant clusters. Shaded areas in c represent the SEM. Cluster-permutation-corrected significant timepoints are indicated with horizontal lines in c. The first part (-0.2-1.2 s) of the time-frequency spectra in a and b and of the time course in c corresponds to the average of short and long trials, and the second part (1.2-3.2 s) corresponds to long trials only. The vertical dotted lines indicate (from left to right) the onset (0 s) and offset (0.2 s) of the retro-cue and the time of probe appearance in short delay trials (1.2 s). PWS: people who stutter; TFS: typically fluent speakers.

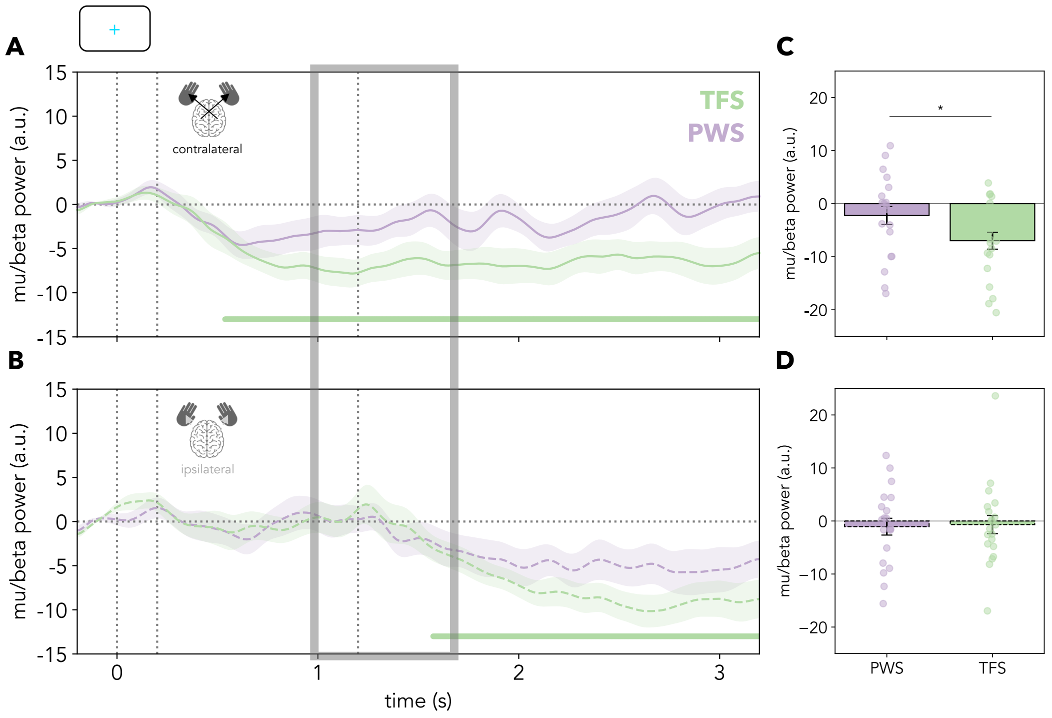


**Supplementary Figure 2. Average mu/beta (8-30 Hz) power in sensorimotor (M1) parcels contralateral and ipsilateral to the cued action, locked cue onset. Related to Figure 2.** (a-b) Mu/beta (8-30 Hz) power relative to the pre-cue baseline (-0.2-0 s) in sensorimotor parcels contralateral (a; solid lines) and ipsilateral (b; dashed lines) to the cued action in TFS (green) and PWS (purple). (c-d) Bar plots of average mu/beta power relative to baseline during the time window displaying a statistically significant difference in lateralised mu/beta modulation between PWS and TFS (1-1.7 s after cue onset; Figure 2c) in sensorimotor parcels contralateral to the cued action (c) and ipsilateral to the cued action (d) in PWS (purple) and TFS (green). Error bars and coloured shaded areas represent the SEM. Gray shaded area in a and b depicts the 1-1.7 s time window used to estimate mu/beta power in c and d. Asterisks represent statistical significance (*$p < .05$; **$p < .01$; ***$p < .001$). The first part (-0.2-1.2 s) of the time courses in a and b corresponds to the average of short and long delay trials, and the second part (1.2-3.2 s) corresponds to long delay trials only. The vertical dotted lines indicate (from left to right) the onset (0 s) and offset (0.2 s) of the retro-cue and the time of probe appearance in short delay trials (1.2 s). Coloured dots represent individual participants. PWS: people who stutter; TFS: typically fluent speakers.

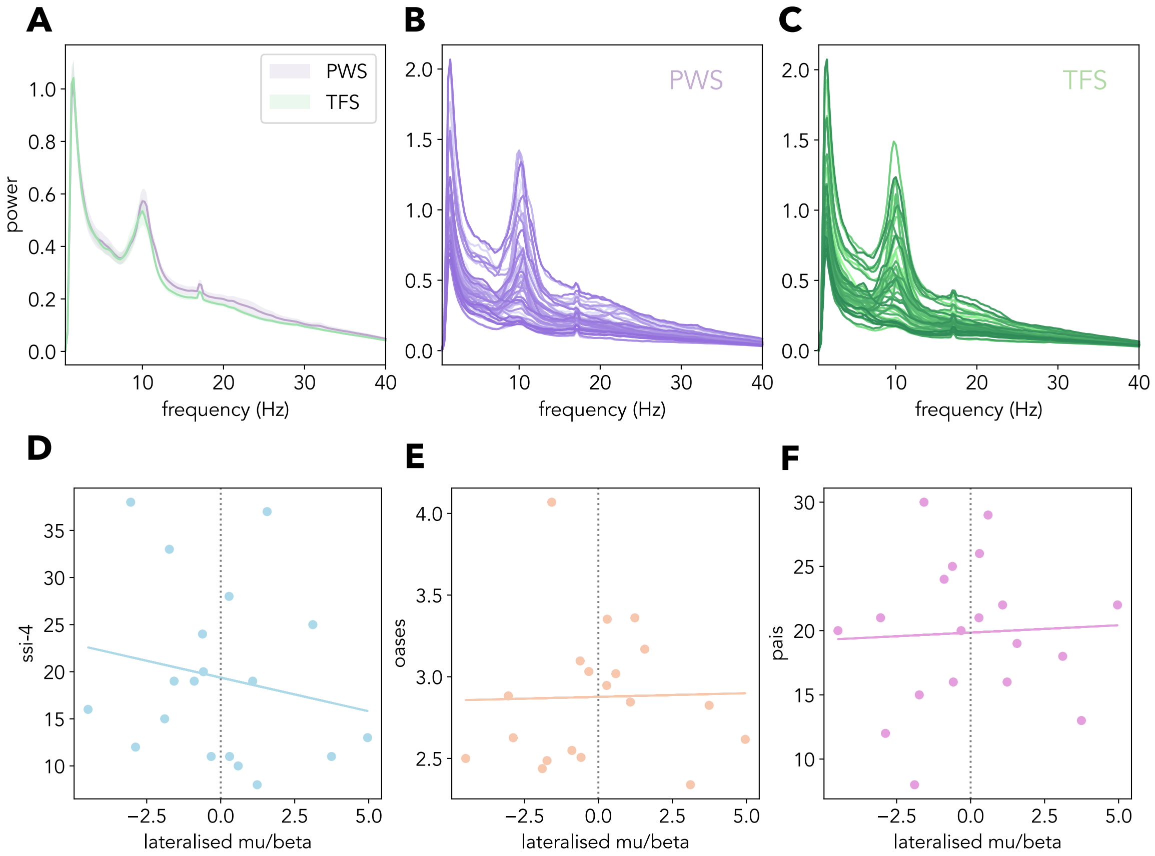


**Supplementary Figure 3**. **Power spectra in PWS and TFS (a-c) and correlations between average lateralised mu/beta modulation (1-1.7 s) and SSI-4 (d), OASES (e), and PAiS (f) scores in PWS (n = 20). Related to Figures 2 and 3.** (a) Parcel-average power spectra in PWS (purple) and TFS (green). (b) Parcel-wise power spectra in PWS. (c) Parcel-wise power spectra in TFS. Across all participants and parcels, an artifact at 17 Hz is observed. (d-f) SSI-4: rho = -.24, p = .31; OASES: rho = .21, p = .4; PAiS: rho = .08, p = .74); SSI-4: Stuttering Severity Index-4; OASES: Overall Assessment of the Speaker's Experience of Stuttering; PAiS: Premonitory Awareness in Stuttering Scale. PWS: people who stutter; TFS: typically fluent speakers.

| **Characteristic** | **TFS (n = 20)** | **PWS (n = 20)** |
|---|---|---|
| **Gender, N (%)** | | |
| Male | 11 (55) | 11 (55) |
| Female | 9 (45) | 9 (45) |
| **Handedness, N (%)** | | |
| Left | 4 (20) | 3 (15) |
| Right | 16 (80) | 17 (85) |
| **Ethnicity, N (%)** | | |
| Asian / Asian British | 5 (25) | 3 (15) |
| Black / Black British / Caribbean / African | 2 (10) | 3 (15) |
| Mixed / Multiple ethnic groups | 2 (10) | 2 (10) |
| White | 11 (55) | 12 (60) |
| Other ethnic group | 0 (0) | 0 (0) |
| **Education level completed, N (%)** | | |
| A-levels / equivalent | 6 (30) | 6 (30) |
| Higher Education / Undergraduate | 8 (40) | 8 (40) |
| Postgraduate / Professional | 6 (30) | 6 (30) |
| **Age, M (SD)** | 24.85 (5.8) | 26.8 (6.29) |
| **Test results, Median [Range]** | | |
| SSI-4 | | 18.5 [11.0-38.0] |
| OASES | | 2.84 [2.34-4.07] |
| PAiS | | 20.0 [8-30] |

**Supplementary Table 1. No differences in gender, handedness, ethnicity, education level and age between typically fluent speakers (TFS; n = 20) and people who stutter (PWS; n = 20). Related to Figure 1 and STAR Methods.**

##### Reaction time (ms)

| group | cue | duration | M | SD |
|---|---|---|---|---|
| PWS | inf | short | 665.93 | 71.83 |
| PWS | noninf | short | 778.71 | 97.19 |
| PWS | inf | long | 527.9 | 69.11 |
| PWS | noninf | long | 678.68 | 64.46 |
| TFS | inf | short | 640.01 | 88.91 |
| TFS | noninf | short | 829.52 | 102.47 |
| TFS | inf | long | 538.39 | 73.15 |
| TFS | noninf | long | 734.98 | 69.74 |

##### log(RT) ANOVA

| factor | df | F value | p value | ges |
|---|---|---|---|---|
| group | 38 | 0.43 | 0.52 | 0.01 |
| cue | 38 | 139.39 | **<.001** | 0.34 |
| duration | 38 | 149.41 | **<.001** | 0.2 |
| group x cue | 38 | 3.53 | 0.07 | 0.01 |
| group x duration | 38 | 1.15 | 0.29 | 0 |
| cue x duration | 38 | 13.43 | **<.001** | 0.02 |
| croup x cue x duration | 38 | 0.29 | 0.59 | 0 |

##### Report error (°)

| group | cue | duration | M | SD |
|---|---|---|---|---|
| PWS | inf | short | 12.11 | 1.51 |
| PWS | noninf | short | 12.55 | 1.46 |
| PWS | inf | long | 13.57 | 1.86 |
| PWS | noninf | long | 13.4 | 2.61 |
| TFS | inf | short | 10.41 | 1.18 |
| TFS | noninf | short | 10.11 | 1.59 |
| TFS | inf | long | 11.32 | 1.23 |
| TFS | noninf | long | 11.94 | 1.8 |

##### log(error) ANOVA

| factor | df | F value | p value | ges |
|---|---|---|---|---|
| group | 38 | 6.52 | **0.02** | 0.08 |
| cue | 38 | 0.75 | 0.39 | 0 |
| duration | 38 | 7.48 | **0.01** | 0.03 |
| group x cue | 38 | 0.13 | 0.72 | 0 |
| group x duration | 38 | 0.25 | 0.62 | 0 |
| cue x duration | 38 | 0.07 | 0.8 | 0 |
| croup x cue x duration | 38 | 0.63 | 0.43 | 0 |

**Supplementary Table 2**. **Summary statistics (left) and ANOVA results (right) of reaction time (ms; top) and report error (°; bottom). Related to Figure 1.** The mean (M) and standard deviation (SD) of reaction time (ms; top left) and report error (°; bottom left) in correct trials for different trial types and groups. ANOVA results for reaction time (top right) and report error (bottom right). Ges (generalised eta squared) reflects the $\eta^2$ metric of effect size. PWS: people who stutter; TFS: typically fluent speakers.

### Linear Mixed-Effects Model

| type | effect | estim. | SE | t | p | var | SD |
|---|---|---|---|---|---|---|---|
| fixed effects | intercept | 0.02 | 0.01 | 2.94 | **0.001** | | |
| fixed effects | group | 0.01 | 0.01 | 0.77 | 0.44 | | |
| fixed effects | within-person amplitude | -0.03 | 0.01 | -2.29 | **0.03** | | |
| fixed effects | between-person amplitude | 0.00 | 0.01 | -0.32 | 0.75 | | |
| fixed effects | within-person rms | 0.02 | 0.01 | 3.02 | **0.001** | | |
| fixed effects | between-person rms | -0.01 | 0.01 | -1.14 | 0.26 | | |
| random effects | intercept | | | | | 0 | 0 |
| random effects | within-person amplitude | | | | | 0.00 | 0.06 |
| random effects | residual | | | | | 0.08 | 0.28 |

**Supplementary Table 3**. **Results of linear mixed effects model. Related to Figure 2.** The coefficients of the fixed and random effects of the LMM that modelled temporal shift ($\tau$) as a function of normalised within- and between-person amplitude and normalised within- and between-person root mean square (RMS; see **STAR Methods**).
